# Sensitive environmental DNA methods for low-risk surveillance of at-risk bumble bees

**DOI:** 10.1101/2025.05.13.649340

**Authors:** Rodney T. Richardson, Grace Avalos, Cameron J. Garland, Regina Trott, Olivia Hager, Mark J. Hepner, Clayton Raines, Karen Goodell

## Abstract

Terrestrial environmental DNA (eDNA) techniques have been proposed as a means of sensitive, non-lethal pollinator monitoring. To date, however, no studies have provided evidence that eDNA methods can achieve detection densities on par with traditional pollinator surveys. Using a large-scale dataset of eDNA and corresponding net surveys, we show that eDNA methods enable sensitive, species-level characterization of whole bumble bee communities, including rare and critically endangered species such as the rusty pathed bumble bee (RPBB; *Bombus affinis*). All species present in netting surveys were detected within eDNA surveys, apart from two rare species in the socially parasitic subgenus *Psithyrus* (cuckoo bumble bees). Further, for rare non-parasitic species, eDNA methods exhibited similar sensitivity relative to traditional netting. Relative to flower eDNA samples, sequenced field negative controls resulted in significantly lower rates of *Bombus* detection, and these detections were likely attributable to high rates of background eDNA on environmental surfaces. Lastly, we found that eDNA-based frequency of detection across replicate surveys was strongly associated with net-based measures of abundance across site visits. We conclude that the method is cost-effective and highly scalable for semi-quantitative characterization of at-risk bumble bee communities, providing a new approach for improving our understanding of species habitat associations.

## Introduction

The global environment is constantly changing, and these fluctuations have important implications for the maintenance of habitable conditions and the availability of natural capital for future generations. The modern era has been marked by accelerated rates of environmental change (Elsen et al., 2022; Mottl et al., 2021; Potapov et al., 2022), making the monitoring of such change an important goal for taxa that provide critical ecosystem services. To better understand changing species distributions, as well as how populations respond to global environmental change, researchers need sensitive and scalable survey methods. This is particularly true for insects, which represent numerous critical taxa, including pollinators, and are notoriously difficult to monitor with traditional methods. With recent advances in the cost-effectiveness and accuracy of DNA sequencing, environmental DNA (eDNA) methods have grown in popularity, with applications now spanning a variety of use cases in both aquatic and terrestrial settings (Allen et al., 2021; Avalos et al., 2024; Jerde et al., 2011; Krehenwinkel et al., 2022). Considering these advances, eDNA methods are expected to complement traditional survey methods by alleviating many of the bottlenecks associated with traditional means of population monitoring, though careful evaluation of performance is needed. Here, we demonstrate the strong potential of terrestrial eDNA methods for large-scale monitoring of bumble bees (genus *Bombus*), a taxonomic group of valuable pollinating species of high conservation concern.

*Bombus* comprise an important component of robust pollinator communities, with over 250 species globally, approximately one third of which are believed to be in decline (Arbetman et al., 2017). Prior to the introduction of European honey bees (*Apis mellifera*), *Bombus* were the only eusocial pollinators in temperate regions of the western hemisphere. With relatively large body sizes and heightened visual capacities (Lunau et al., 2009), these species exhibit unique traits relative to most Anthophila (bees), including the ability to forage in low-light conditions, at low temperatures and over long-distances (Goulson, 2003; Heinrich, 2004; Tichit et al., 2024). Such traits make *Bombus* ideal for pollination in cold climates, forest understories and on plant species that require buzz pollination, including numerous agriculturally important crops (Kendall et al., 2019; Miller-Struttmann et al., 2017; Oyen et al., 2016). Thus, the services provided by *Bombus* are often difficult to replace with other pollinator taxa (Brosi & Briggs, 2013; Willmer et al., 1994). Unfortunately, *Bombus* have experienced range contractions and widespread turnover in community composition over recent decades, largely driven by dramatic declines in formerly widespread species. (Arbetman et al., 2017; Grixti et al., 2009; Morales et al., 2013). These changes raise concern about the future of this functionally important pollinator group.

In North America, there is strong evidence of declines within the subgenera *Bombus* and *Thoracobombus* (Cameron et al., 2011; Colla & Packer, 2008; Graves et al., 2020). In 2017, the rusty patched bumble bee (RPBB; *Bombus affinis*) was listed under the US Endangered Species Act (Christopher, 2017). At the time of listing, the species was widely believed to be restricted to areas of the Upper Midwest US until recent work revealed a persistent population within the Central Appalachian mountains (Hepner et al., 2024). Accordingly, the Central Appalachian Mountains are an under-surveyed region of high conservation potential that also harbors populations of other bumble bee species of concern, including *B. terricola* and *B. pensylvanicus*. To improve our knowledge of *Bombus* communities within this region, and test novel detection methods, we conducted a large-scale survey of the region using both traditional netting and novel environmental DNA methods. Here, we evaluate the general efficacy of our eDNA methods and compare them to netting in terms of detection sensitivity, taxonomic breadth of detection, and quantitative reliability.

## Methods

From June to September 2022, we conducted 84 site visits across 55 locations. Sites consisted of linear roadside rights-of-way, open areas or forest understories with abundant floral resources, typically 0.25 to 0.5 hectares in area. The sampling region predominately included the Central Appalachian Mountains, with one additional site near Easton, Maryland (MD). Survey regions were selected based on permitted access to public and some private lands based on local, state and federal permits. Within these regions, specific sites for surveys were selected based on the availability of flowers attractive to bumble bees. We typically stayed close to navigable roads to maximize the area covered and number of survey locations. During each site visit, eDNA and netting surveys were conducted in series, with 2 – 5 eDNA and 1 – 6 net surveys completed. When flower patches were dense but only encompassed a small spatial area, multiple eDNA samples could be collected efficiently, while the feasibility of repeating net samples was limited. When surveying sites of perceived high-value (i.e., dense patches of bumble bee-attractive flowers) that encompassed large spatial areas, we increased net survey intensity relative to eDNA sampling. Notably, samples collected for this work have been reported on previously in an unrelated manuscript on the use of eDNA methods for broad-scope surveillance of all Arthropoda (Richardson et al., 2025).

### eDNA Sample Collection

During each site visit, flower eDNA and leaf surface eDNA samples were collected. Leaf surface samples were intended to represent field negative controls and included leaves from trees or shrubs that were at least 3 m from prominent floral forage, and which showed no evidence of insect herbivory or visible debris. For each eDNA sample, flowers or leaves of a single plant species were clipped into a sterile quart-sized, zip-closable plastic bag until the bag was approximately ¼ to ¾ full of loose plant material. Samples were mixed with 25 mL of eDNA preservative (10 percent EtOH v/v, 40 percent propylene glycol v/v and 0.25 percent sodium dodecyl sulfate w/v), mixed gently and allowed to sit for 1 to 2 min before the eDNA-containing preservative rinse was carefully poured into a labelled 50 mL conical vial for transport to the laboratory. During eDNA collection, researchers used flame-sterilized tools and wore disposable sterile nitrile gloves to avoid cross-contamination between sites or between netting and eDNA surveys. To further reduce the risk of contamination between methods, sampling of eDNA was conducted prior to corresponding net surveys.

### Traditional Netting Surveys

During each 10-min net survey, an observer walked a meandering transect, independent of other observers, through areas with dense floral cover while searching for bees. For most surveys, observers netted continuously and indiscriminately, targeting bumble bees while also opportunistically collecting other bee species. Following these ‘indiscriminate surveys’, live bees were transferred to a one gallon zip-closable bag using the technique described in Hepner, (2024). Any *B. affinis* present in the sample was transferred to a clean glass vial after carefully making an incision in the bag, chilled briefly on ice, then photo-documented and released. Other species of conservation concern, including *B. pensylvanicus* and *B. terricola*, were also often removed from the sample and released to limit lethal take of these species to one worker or male per site visit. During site visits where the perceived risk of netting *B. affinis* was high, we modified our survey methods to reduce the risk of harm to individual bees. In these situations, observers conducted ‘targeted visual’ surveys, where bumble bees were netted and handled individually and observers focused exclusively on a suite of high conservation value species that are distinguishable in the field, including *B. affinis, B. pensylvanicus, B. terricola, B. fervidus*, and *B. auricomus*. Data from these targeted visual surveys were only used for certain analyses, as described below, since they were not sensitive for detection of *B impatiens, B bimaculatus, B griseocollis, B perplexus, B. vagans, B. sandersoni, B. flavidus* and *B. citrinus*. Bumble bee specimens that were lethally collected were pinned using standard entomological practices or frozen individually in 1.5 mL microcentrifuge tubes. We identified each specimen using microscopic examination of characters detailed in Colla et al. (2011), and Williams et al. (2014). Specimens of cryptic species, *B. perplexus, B. vagans* and *B. sandersoni*, were distinguished with a combination of characters from Milam et al. (2020) and inner hind tibial spur differences (Z. Portman, University of Minnesota, *pers. comm*.).

### Laboratory eDNA Processing

A custom salt-ethanol precipitation protocol was used to pellet DNA from each sample and follow-up purification procedures, including use of the Qiagen PowerClean Pro Cleanup Kit, were used for further removal of PCR inhibitors. Readers can find full details of this process in Richardson et al. (2025). Following DNA isolation, Illumina MiSeq amplicon libraries were prepared using a 3-step PCR-based protocol established in previous work (Richardson et al., 2019), which relies on slight modifications of previously established methods (Kozich et al., 2013). During library preparation, the bumble bee-specific COI primers from Milam et al. (2020) were used for the initial amplification. To quantify critical-mistagging rates during downstream sequencing (Esling et al., 2015), we incorporated 15 no-library negative control dual-index combinations during library preparation. Amplified libraries were purified with the SequalPrep Normalization Plate Kit, pooled equimolarly and subjected to Illumina MiSeq sequencing using a 2 x 150 cycle v2 flow cell.

### Taxonomic Annotation of eDNA Data

After sequencing, VSEARCH v2.8.1 (Rognes et al., 2016) was used to merge paired-end COI sequences and remove priming sites from the 5’ and 3’ ends. Sequences were taxonomically annotated by semi-global VSEARCH top-hit alignment against a custom reference sequence database trimmed to the amplicon regions of interest using MetaCurator (Richardson et al., 2020), with dependencies HMMER v3.1b2 (Eddy, 2011), MAFFT v7.270 (Katoh et al., 2002), VSEARCH and Taxonomizr v0.11.1 (Sherrill-Mix, 2019). During alignment, a minimum query cover of 0.8 was required. The reference database was constructed using sequences from Eastern North American *Bombus* available through NCBI, downloaded on February 2^nd^, 2023. Following reference data curation, the database consisted of 61 sequences representing all species in the sampling region except the parasitic *B. variabilis* and *B. insularis*. Notably, all species were represented by two or more reference sequences with the exception of *B. fraternus*. During alignment, 100 percent identity matches were considered confident species-level detections, and this threshold was strongly supported by leave-one-out cross-validation analysis of available reference data (Supplemental Figure S1). Since all Illumina sequencing runs produce low frequency misidentifications when inferring dual-index tags, leading to critical mistags (Esling et al., 2015), we used an established technique to remove identifications with greater than 0.005 probability of representing a critical mistag-associated detection (Richardson, 2022). Additionally, we removed any detections which were represented by two or fewer sequences in a sample. All computational analysis was performed on the Owens cluster of the (*Ohio Supercomputer Center*, 1987).

### Statistical Analysis

Statistical analysis of all resulting data was conducted using R (R Core Team, 2021). Following taxonomic annotation of sequences, leaf surface eDNA samples were compared against flower eDNA samples using a *X*^*2*^ test to evaluate differences in frequency of eDNA detection per sample and a Wilcoxon Rank Sum test to evaluate differences in *Bombus* species richness per eDNA sample. To analyze species detectability and occupancy patterns, the spOccupancy package (Doser et al., 2022) was used to produce an integrated occupancy model for each species in the dataset. Integrated occupancy models allow for the specification of multiple detection processes, eDNA and netting in this case, each of which informs occupancy estimation. Across methods, significant differences in per species detection or occupancy was assessed based on Bayesian 95 percent credible interval overlap. Since netting and eDNA data were collected in a relatively paired fashion and our main goal was to compare these survey methods, null intercept-only models were specified for both the occupancy and detectability components of each model. To broadly compare detection sensitivity across methods, we regressed eDNA detectability estimates against net survey detectability estimates for each species using ordinary least squares regression. Lastly, on a per site visit basis, we regressed the eDNA-based frequency of detection of each species against the log-transformed mean number of *Bombus* individuals per net survey to assess the degree to which eDNA detection predicts species abundance across sites. A binomial generalized linear mixed-effects model (glmmTMB; Brooks et al., 2017) was used for this analysis, with species and sampling site specified as crossed random intercept terms to account for repeated measures. Regressions were performed separately for the flower and field negative control eDNA samples and we only used data from indiscriminate surveys for this analysis since targeted visual surveys provided no sensitivity for detecting 5 of 13 species present in the study system.

## Results

Similar numbers of samples were taken using each survey method (Figure 1A), with a total of 251 flower eDNA samples, 22 field negative control eDNA samples, 165 indiscriminate 10-min net surveys and 103 targeted visual surveys. Across the study, we detected 13 *Bombus* species, and the frequencies of detection for each species were similar between the two survey methods (Figure 1B).

**Figure 1.**
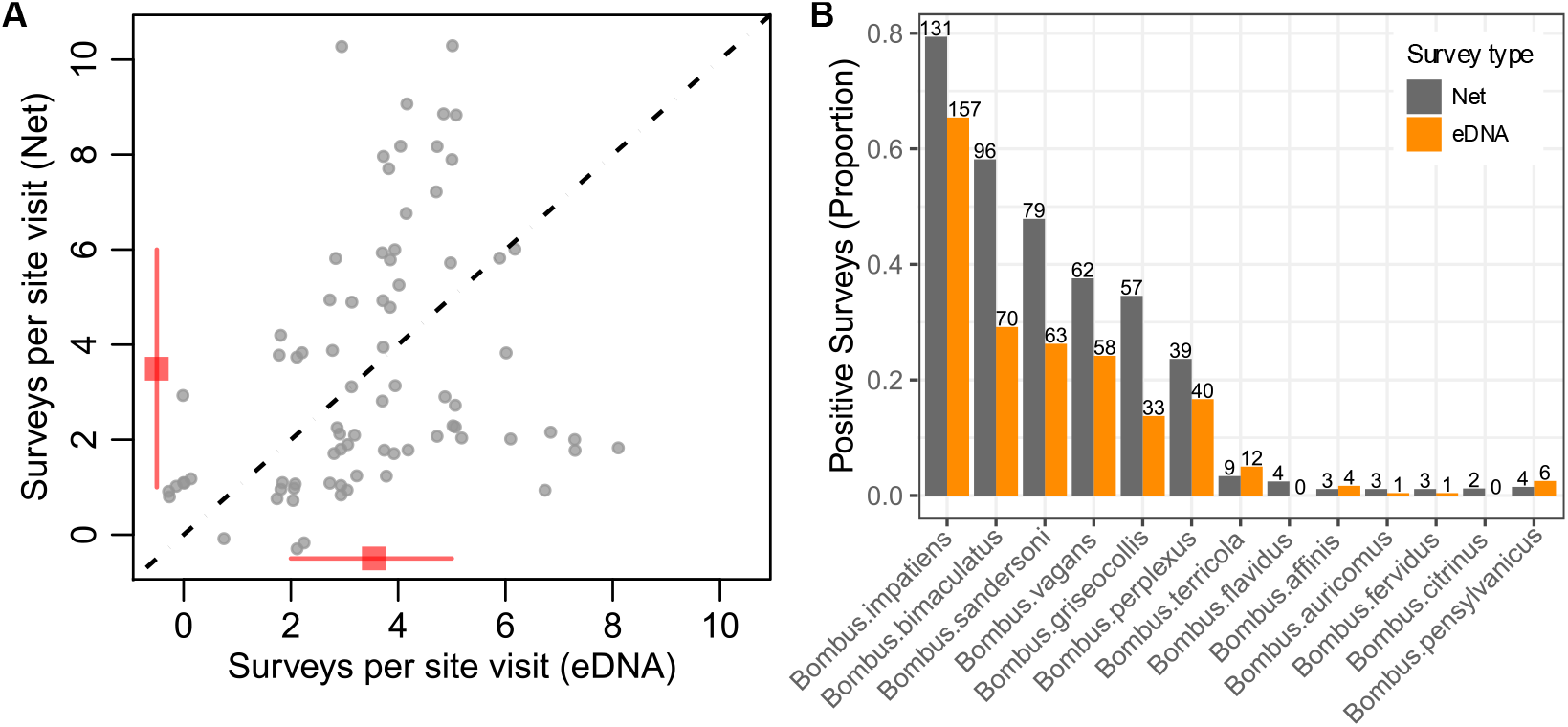
Pairwise comparison of sampling effort per site visit for each survey method (A), where points were jittered using random draws from a *U*(-0.3, 0.3) distribution and the mean, 20^th^ and 80^th^ quantiles are shown with square points and lines. A bar plot (B) shows comparison of frequency of *Bombus* species detection, with numbers above bars indicating the count of positive surveys for each method. For eDNA bars, data from N=251 flower eDNA samples were included and field negative controls were excluded. For netting bars, data from N=268 10-minute net surveys (165 indiscriminate net surveys and 103 targeted visual surveys) were used to estimate frequency of detection for species of conservation concern, listed in the methods, while N=165 indiscriminate net surveys were used for the remaining common species.

### eDNA Survey Results

Among flower eDNA surveys, sampled plant species spanned 20 families, with the bulk of samples originating from species within Asteraceae (N = 109), Lamiaceae (N = 49) and Fabaceae (N = 43). Illumina sequencing yielded a total of 5.7 million successfully mate-paired reads, with a median sequencing depth of 20,850 sequences produced per flower eDNA sample. Field negative control eDNA samples yielded significantly fewer sequences, with a median of 528 sequences per sample (Figure 2A, Wilcoxon rank-sum test: *P* = 0.006). Most sequences, 63 percent in total, belonged to bumble bees and there was minimal evidence of variance in this percentage across flower and field negative samples (Figure 2B, Wilcoxon rank-sum test: *P* = 0.086). As expected, flower eDNA samples exhibited significantly greater frequency of *Bombus* detection (Figure 2C; *X*^*2*^ = 8.89, *P* = 0.001) and significantly greater *Bombus* species richness (Figure 2D, Wilcoxon rank-sum test: *P* < 0.001) relative to field negative controls. Out of the 13 *Bombus* species recorded across all surveys, eDNA methods detected 11, including multiple detections of the federally endangered *B. affinis*. Both species that went undetected using eDNA belonged to the socially parasitic sub-genus *Psithyrus*. Among 22 field negative control eDNA samples, a total of 15 *Bombus* detections were observed. Negative control detections were composed predominately of highly abundant species including *B. impatiens* (N = 6), *B. bimaculatus* (N = 4) and *B sandersoni* (N = 3), with one detection each for *B. vagans* and *B griseocollis*.

**Figure 2.**
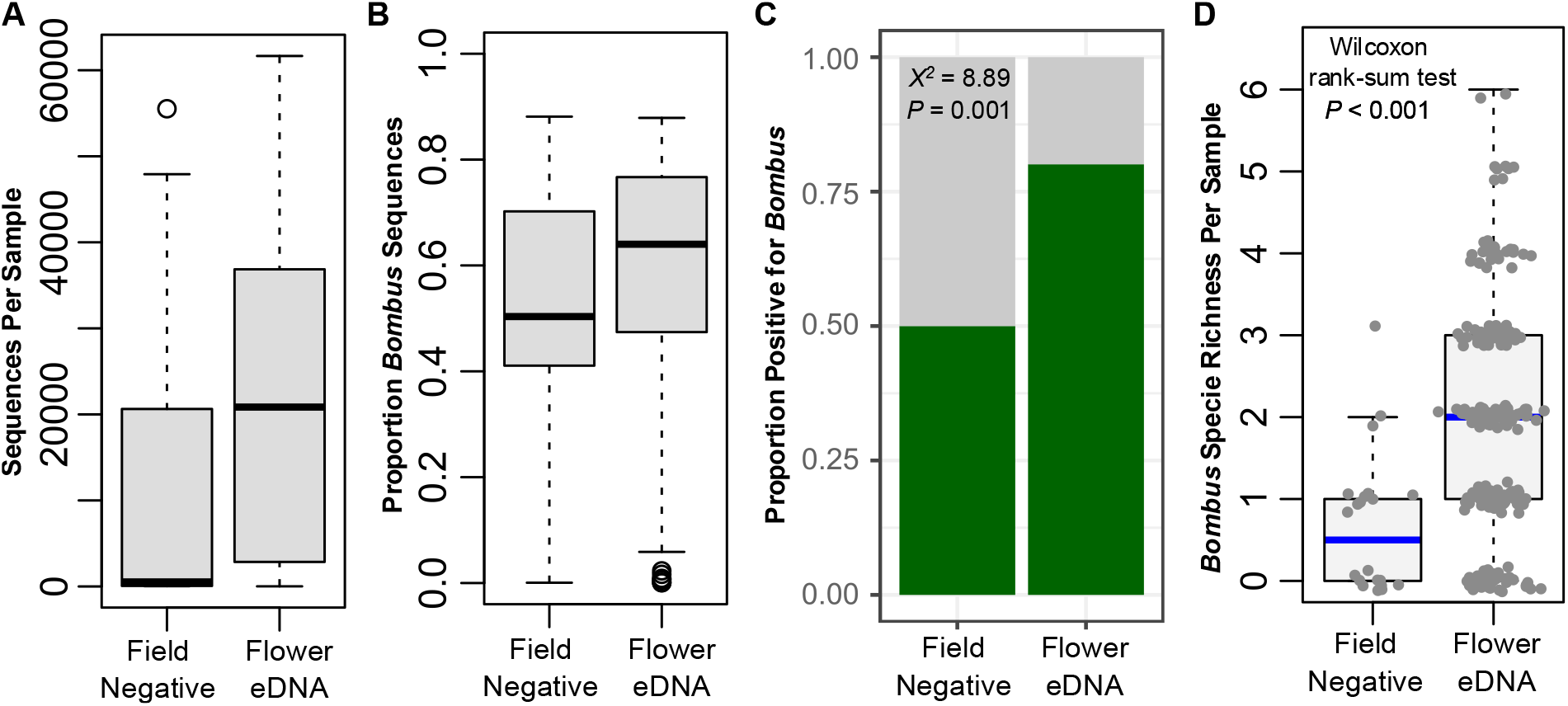
Comparison of flower (N = 251) and field negative control (N = 22) eDNA samples, in terms of total sequences per sample (A), proportion of sequences assigned to *Bombus* per sample (B), frequency of *Bombus* detection (C) and *Bombus* species richness per sample (D).

### Net Survey Results

Net surveys resulted in a total of 2,769 lethally collected *Bombus* specimens from indiscriminate surveys, as well as non-lethal observation of approximately 2,074 additional individuals during targeted visual surveys. Notably, targeted visual surveys yielded 12 observations of species of conservation concern, including 1 observation of *B. terricola* and 11 observations of *B. pensylvanicus* (all at a single site near Easton, MD). Among lethally collected specimens, 2,759 were identified to species. The remaining 10 specimens could not be identified due to the poor condition of diagnostic features.

### Comparison of eDNA and Net Results

Integrated occupancy modelling of each species resulted in well-converged MCMC models (maximum Gelman-Rubin 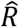= 1.04) with large effective sample sizes (minimum ESS = 1,059). Model results revealed similar detection probabilities between eDNA and netting (Figure 3A), with some notable deviations. Overall, per species mean detection probabilities were highly correlated across the two survey methods (OLS Regression, *P* = 0.001, *R*^*2*^ = 0.60). However, for species within the subgenus *Pyrobombus*, eDNA displayed significantly lower detection probabilities relative to netting for 4 of 5 species. Similarly, eDNA detection sensitivity was significantly less than that of netting for the only species of *Cullumanobombus* in the data, *B. griseocollis*. While eDNA exhibited relatively lower detection sensitivities for species within the subgenera *Psithyrus* and *Bombias*, sample sizes were minimal, obscuring any assessment of significance. For species within the subgenera *Bombus* and *Thoracobombus*, detection sensitivities were similar across the two methods, with no significant differences. Occupancy estimates across the sampled sites varied considerably among species (Figure 3B), with means ranging from 0.03 (*B. pensylvanicus*) to 0.98 (*B. impatiens*). Mean occupancy of *B. affinis* was 0.22, with lower and upper credible bounds of 0.07 and 0.59, respectively. For species of primary conservation concern, *B. affinis* and *B. terricola*, eDNA and net-based observations occurred within heavily forested montane landscapes (Figure 5A and 5B), consistent with the analysis presented in Hepner et al. (2024). A third species of heightened concern, *B. pensylvanicus*, was observed at only a single site (Figure 5C), though this species is known to be associated with grasslands (Novotny et al., 2021), which were relatively rare in the landscapes we sampled. Lastly, investigation of the relationship between eDNA-based frequency of detection and net-based species abundance revealed highly significant associations for both flower eDNA (Figure 4A: *P* < 0.001) and field negative control eDNA (Figure 4B: *P* = 0.008) samples.

**Figure 3.**
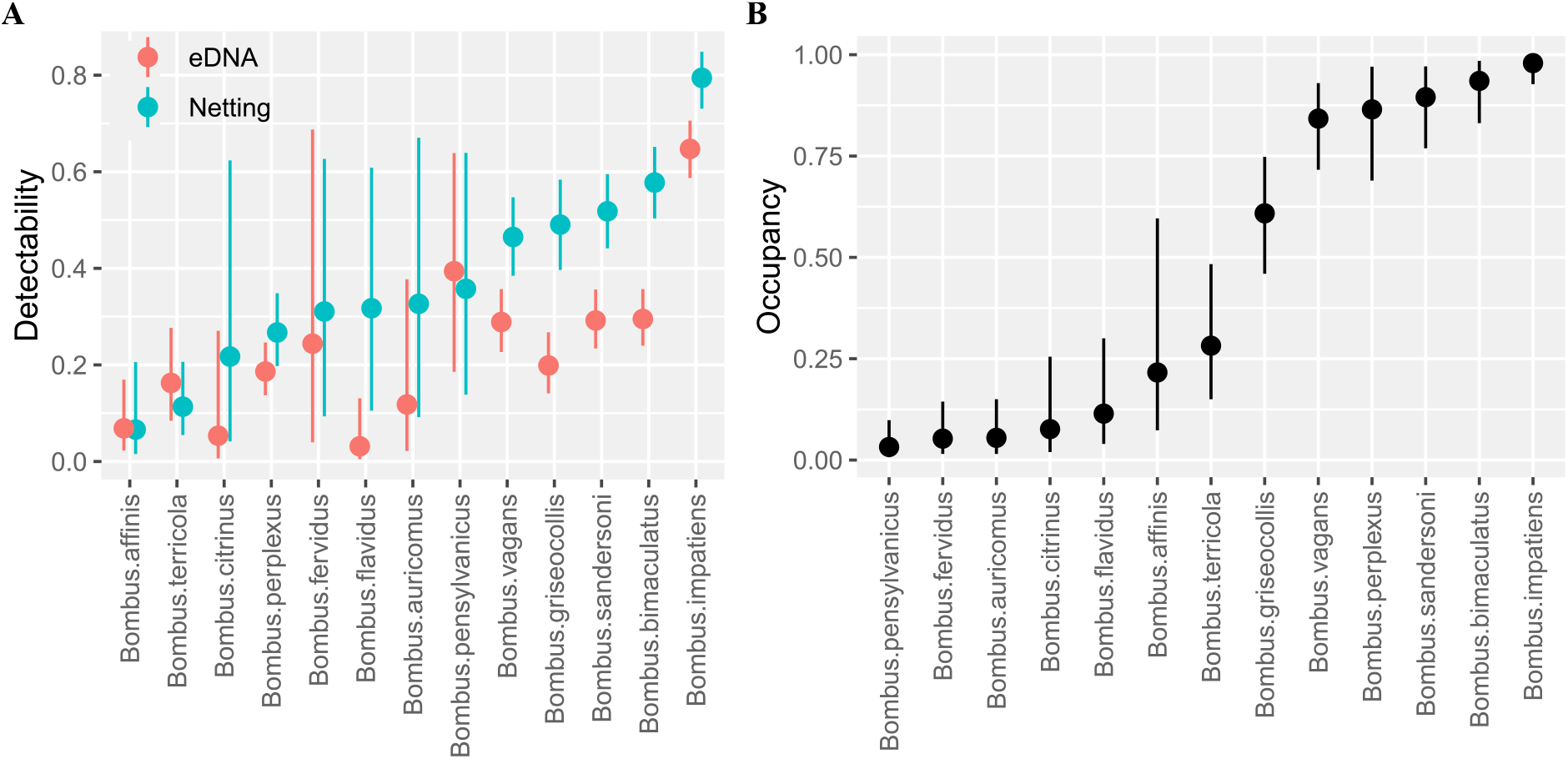
Comparison of species detection probabilities between eDNA and netting (A) and estimates of species occupancy across the survey sites (B). Occupancy estimates were inferred using both eDNA and net detections with integrated single-species models (Doser et al., 2022). Points indicate means, with lines extending to 95 percent credible bounds. For eDNA, field negative control results were excluded and inferences are from N=251 flower eDNA samples. For species of conservation concern (list provided in Methods), netting inferences are from N=165 indiscriminate net surveys and N=103 targeted visual surveys. For common species, netting inferences are from N=165 indiscriminate surveys since there was no sensitivity for common species within targeted visual surveys.

**Figure 4.**
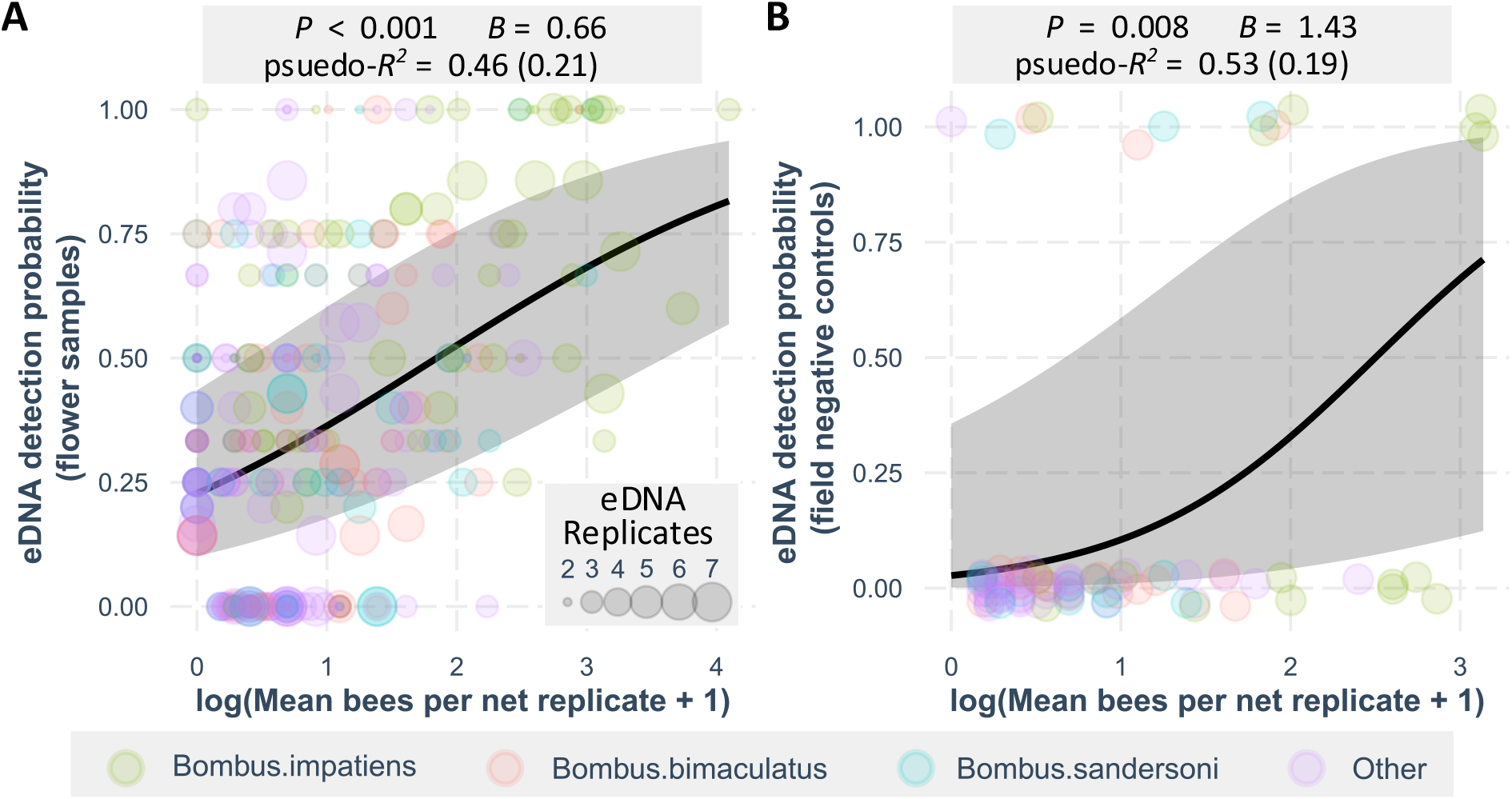
Relationship between eDNA-based frequency of detection and log-transformed mean number of *Bombus* individuals per net survey for flower eDNA samples (A) and field negative control eDNA samples (B). Each point represents a unique site visit and, for flower samples, point sizes represent the number of eDNA samples collected during the visit. Conditional Nakagawa & Schielzeth (2013) *pseudo-R*^*2*^ values are provided, with marginal estimates shown in parentheses.

**Figure 5.**
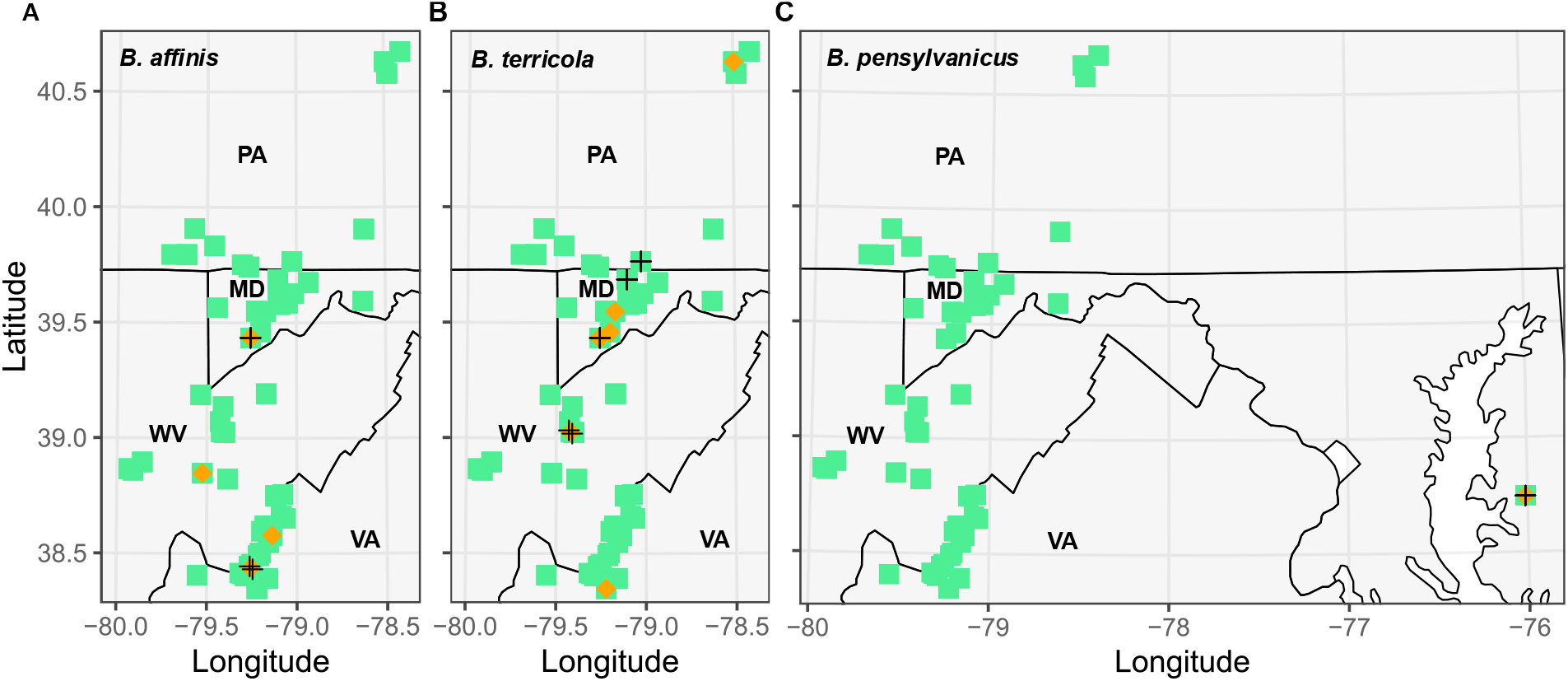
Detection locations for primary species of conservation concern, *B. affinis* (A), *B. terricola* (B) and *B. pensylvanicus* (C). Spatial breadth of sampling is shown in green squares, with eDNA detections shown as orange diamonds and net detections represented by black crosses. To aid visualization, the coastal Maryland site was excluded for cases in which the species of interest was not detected.

## Discussion

Widespread environmental change, coincident with mounting evidence of declining wildlife populations, necessitates increased monitoring to support conservation management. But limited scientific funding and cumbersome, often risky, survey methods present a considerable challenge for the monitoring of endangered species (Guénard et al., 2025; McCarthy et al., 2012). Within pollinator research, reliance on traditional techniques often limits sample size and spatial replication (e.g. Boone et al., 2023; Nunes et al., 2024; Otto et al., 2023). Efforts to integrate diverse datasets within analyses can increase sample sizes (Ellis et al., 2025; Janousek et al., 2023; Hepner, 2024), but only at the cost of meaningful drawbacks. Integrating data from numerous sources that change over time inevitably introduces systematic bias regarding how, where, by whom, and for what purpose, samples were collected. For pollinators in particular, Portman (2022) provides an excellent demonstration of how such analyses can be strongly confounded. With these trade-offs, researchers consistently express need for improved monitoring techniques (Rousseau et al., 2024).

Environmental DNA methods are increasingly cost-effective, highly scalable and easily distributed across spatially dispersed teams of scientists. Highly replicated survey designs can be implemented by minimally trained personnel, minimizing risk to target species of conservation concern while maximizing the precision with which detection bias can be quantified. For protected species, the permitting process for eDNA surveys is easier relative to more invasive and risky approaches involving handling of individuals. In some cases, including for surveys of reproductive queens of federally protected *Bombus* in spring and fall, eDNA is likely the only permissible survey approach available. eDNA methods are also highly amenable to probability sampling, an approach that limits bias in population monitoring (Boyd et al., 2023), but has not been used in pollinator monitoring to our knowledge.

In comparing eDNA methods against traditional surveys, eDNA and net survey data revealed highly similar inferences of the regional bumble bee community at both regional and local scales. The similarity was true both qualitatively and quantitatively, suggesting that eDNA and traditional methods yield similar inferences regarding *Bombus* species presence and relative abundance at the site level. To our knowledge, these results are unprecedented for a pollinator-oriented study. Past pollinator eDNA studies, including our own, have generally struggled to demonstrate detection sensitivity and taxonomic breadth of detection on par with traditional surveys (Avalos et al., 2024; Gamonal Gomez et al., 2023; Kestel et al., 2023; Newton et al., 2023; Stothut et al., 2024).

In addition to establishing a new technique for *Bombus* surveillance, we provide baseline estimates of occupancy within the Central Appalachian region. We find that while the federally endangered RPBB is present across much of the sampling region, with detections in Virginia, Maryland, and West Virginia, it is difficult to detect. RPBB detection probabilities were among the lowest of all 13 species detected for both eDNA and netting. In concert with a limited number of total detections, modelling suggested a high degree of uncertainty in regional RPBB occupancy (Figure 3). The closely related yellow-banded bumble bee (YBBB; *B. terricola*), which has suffered declines similarly to RPBB (Cameron et al., 2011; Grixti et al., 2009; Jacobson et al., 2018), exhibited similar estimates for detectability and occupancy in our study. Interestingly, five species exhibited mean occupancy estimates which were less than those of RPBB or YBBB. These included two parasitic species from the subgenus *Psithyrus* as well as three species considered to be grassland associates: *B. auricomus, B. fervidus* and *B*. pensylvanicus (Novotny et al. 2021). The low levels of occupancy observed for these species may reflect natural life history traits more so than current conservation status within the region. For grassland-associated species, occupancy would be expected to be low since sampling locations were heavily forested, with a median forest cover of 87.8 percent at a 3-km radius scale. Regarding parasitic *Psithyrus*, species within this group are unique from other *Bombus* in that they only exist as queens and males with no production of workers. Further, *Psithyrus* do not engage in foraging to support colony growth and reproduction, instead foraging only on nectar to meet individual metabolic needs (Bower et al., 2023; Lhomme & Hines, 2019). Altogether, without any prior regional estimates of occupancy or abundance for these groups in the past, it is difficult to assess current status. Accordingly, our results provide baseline estimates for comparison with future *Bombus* monitoring outcomes.

Regarding eDNA quality control, the robustness of our work was supported by the inclusion of multiple forms of controls within our field collection and laboratory processing. Leaf surface samples were included for each site visit as a form of field negative control, and we observed significantly lower frequency and richness of *Bombus* detection within these samples. Of the *Bombus* detections that occurred in leaf surface samples, some may represent cross-contamination within the field or lab. However, increasing evidence suggests that eDNA disperses broadly within the environment (Allen et al., 2023; Valentin et al., 2021), likely facilitated by airborne dispersal (Bohmann & Lynggaard, 2023; Garrett et al., 2023; Johnson, Barnes, et al., 2023). In a companion insect eDNA study of the samples analyzed here, frequency and richness of detections were statistically indistinguishable across leaf and flower eDNA samples for Lepidoptera and all Arthropoda (Richardson et al., 2025), suggesting that *Bombus* detections within leaf samples plausibly represent genuine eDNA signal. Accordingly, a number of works have emphasized the potential of eDNA to illuminate plant-pollinator associations (Avalos et al., 2024; Johnson, Katz, et al., 2023), but our work suggests researchers should be cautious about over-interpreting the eDNA data. Such studies will likely require the inclusion of large numbers of carefully selected background eDNA samples to appropriately adjust for rates of genuine background detection.

In our work, only a single technical replicate was processed for each sample and sample sequencing depths were modest relative to past works (see Table 1, Richardson et al., 2025). Current eDNA sensitivity for *Bombus* appears competitive with traditional approaches, and there is considerable potential to improve this sensitivity going forward. Achieving greater statistical power during eDNA surveys can be obtained in the field stage through additional biological replication, or in the laboratory analysis stage by increasing technical replication or sequencing depth. Approaches to improving the statistical power for traditional survey techniques are more limited and can only be accomplished during field work, where there is little opportunity for cost-savings. Accordingly, we believe our approach will be highly applicable to both near and long-term pollinator conservation efforts, serving as a template for development of terrestrial eDNA survey methods for other rare or cryptic taxa.

## Supporting Data

All sequence data and relevant summary tables needed to reproduce these analyses are publicly available at https://doi.org/10.5281/zenodo.16985008.

## Acknowledgements

This work was primarily supported by a DoD ESTCP Grant to RTR and KG (Project RC22-B5-7373). Additional support was provided by the Maryland Department of Natural Resources Power Plant Research Program and Western EcoSystems Technology, Inc.’s Wildlife Research Initiative; the Appalachian Laboratory of the University of Maryland Center for Environmental Science; and Metamorphic Ecological Research and Consulting, LLC. For land access, permitting and coordination, we thank USFWS, USFS, George Washington and Jefferson National Forest, VA DCR-DNH, MD Forest Service, MD Park Service, MD DNR, WV DNR, WV State Parks, PA Game Commission and Easton Utilities. Any use of trade, firm, or product names is for descriptive purposes only and does not imply endorsement by the U.S. Government.

## Supplementals

**Supplemental Figure S1.**
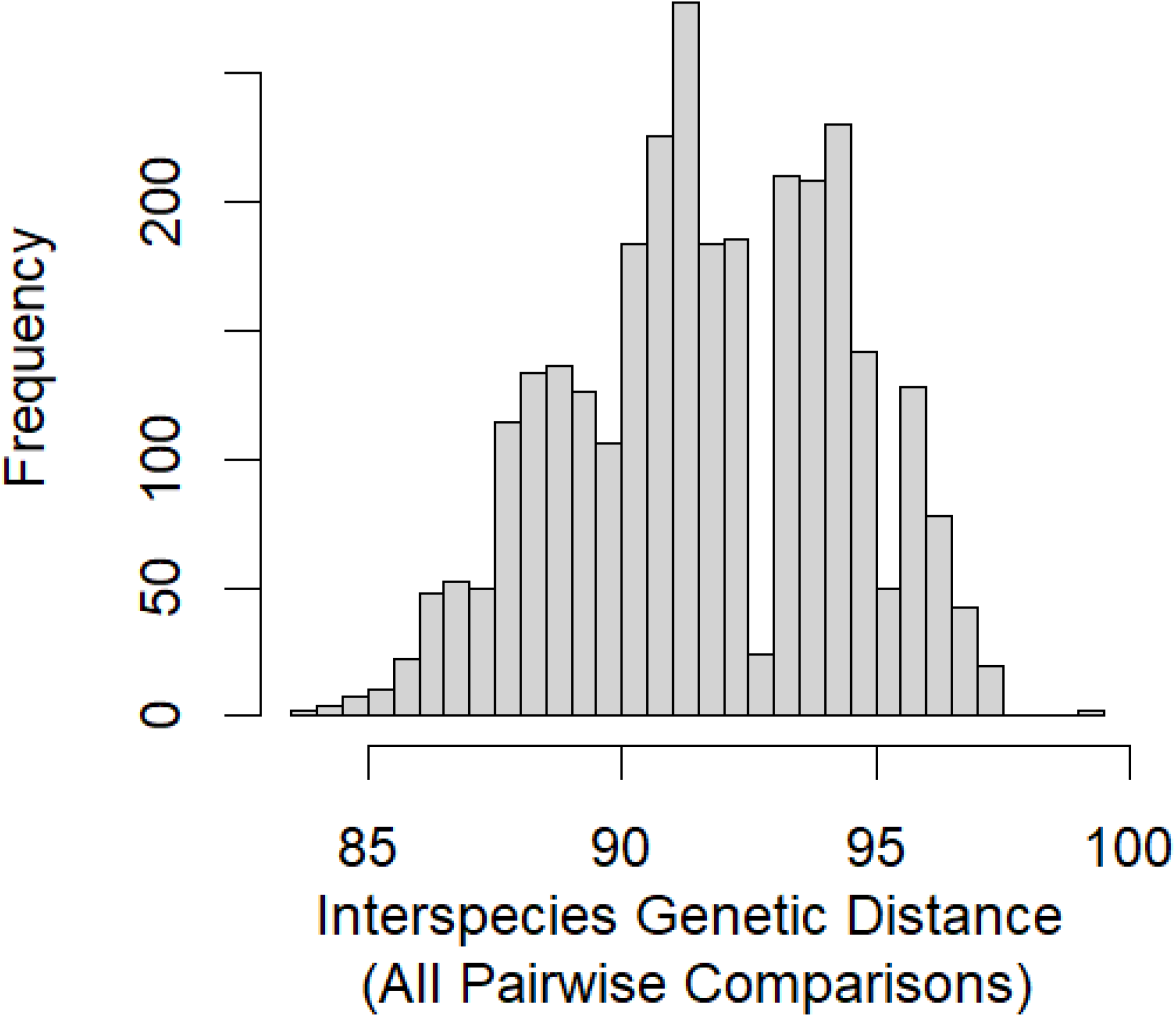
Interspecies genetic distances of all pairwise comparisons of available Eastern North American *Bombus* reference sequence data. Two outlier sequences were 99.4 percent identical to one another; however, these sequences belonged to *B. vagans* and *B sandersoni*, species which are notably difficult to distinguish from one another (Milam et al. 2020). All other pairwise comparisons resulted in alignment mismatches of 2.5 percent, or more, equivalent to 3 or more mismatched base pairs for this genetic marker. Since 100 percent matches were required for species annotation, the probability of having 3 mismatches with locations and base pair substitutions corresponding exactly to a reference is expected to be extremely low.

